# MDM-TASK-web: MD-TASK and MODE-TASK web server for analyzing protein dynamics

**DOI:** 10.1101/2021.01.29.428734

**Authors:** Olivier Sheik Amamuddy, Michael Glenister, Özlem Tastan Bishop

## Abstract

MDM-TASK-web is the web server for the MD-TASK and MODE-TASK software suites. It simplifies the set-up required to perform and visualize results from dynamic residue network analysis, perturbation-response scanning, dynamic cross-correlation, essential dynamics and normal mode analysis. In a nutshell, the server gives access to updated versions of the tool suites, and offers new functionalities and integrated 2D/3D visualization. An embedded work-flow, integrated documentation and visualization tools shortens the number of steps to follow, starting from calculations to result visualization. The web server (available at https://mdmtaskweb.rubi.ru.ac.za/) is powered by Django and a MySQL database, and is compatible with all major web browsers. All scripts implemented in the web platform are freely available at https://github.com/RUBi-ZA/MD-TASK/tree/mdm-task-web and https://github.com/RUBi-ZA/MODE-TASK/tree/mdm-task-web.

**Highlights:**

- MDM-TASK-web is the web server for highly utilized MD-TASK and MODE-TASK with updates
- Eight residue network centrality metrics are available to analyze static and dynamic proteins
- Novel comparative essential dynamics is established to compare independent MD simulations
- Communication propensity tool to evaluate residue communication efficiency is implemented.
- Normal mode analysis from static and protein MD simulations is provided

## 1. Introduction

Molecular dynamics (MD) simulations are a very useful method of conformational sampling to study the dynamics of proteins. Due to large number of internal degrees of freedom for atomic motion, and the complex interactions found within biological macromolecules, investigating the effect of local differences within the protein systems using conventional metrics such as root mean square deviation (RMSD), root mean square fluctuation (RMSF) or the radius of gyration (Rg) may not be sufficient. Network analysis has the ability to abstract out such complexity while maintaining the inter-residue relationships, from which several centrality metrics derived from the social sciences, may be applied to investigate protein dynamics. Many research groups have applied residue interaction network (RIN) analysis on static structures, and have used multiple strategies and tools for summarizing the protein interactions using various edge and node modeling approaches [1–4] to minimize bias and maintain enough variance for recapitulating proteins topological changes. We previously proposed a post-hoc analysis approach of MD simulations using dynamic residue network (DRN) analysis to probe the impact of mutations [5,6] and allosteric effects [7], before setting up the MD-TASK tool suite [8] in 2017. The tool introduced the concept of averaging residue network metrics over MD simulations, as an alternative to examining single networks, to consider the dynamic nature of functional proteins.

Timescales for observing functionally-relevant changes vary with different orders of magnitude, and can also be manifested within very large biological assemblies such as viral capsids, which can make thorough modeling of such systems computationally expensive. Normal mode analysis (NMA) and essential dynamics are two useful methods to study protein dynamics, and their structural and functional behavior [9–11]. Our second software suite, MODE-TASK [12] was developed for the analysis of large-scale protein motions, and comprises ANM from static protein structures, and essential dynamics (ED).

Both MD-TASK and MODE-TASK have been highly utilized to mine protein dynamics by offering a series of novel approaches that have demonstrated their applicability in a growing number of cases [13–21]. Although both software suites are relatively easy to use, the required technical knowledge and software dependencies may act as a hurdle against more widespread usage of the tools and techniques, due to the diversity of operating systems and the relatively fast-evolving Python libraries. We aimed to bridge this gap by providing access to both tools while introducing new functionalities in MDM-TASK-web. It has been designed with a simple and intuitive interface that is supported by any recent web browser. The need for additional software, complex dependencies and command line expertise is greatly reduced. MDM-TASK-web includes new features such as additional network centrality metrics for both DRN and Residue Interaction Network (RIN) centrality calculations from single structures, a communication propensity (CP) tool [22,23], an aggregator of weighted residue contact maps, comparative ED, an anisotropic network model (ANM) workflow, normal mode analysis from MD and integrated 2D/3D visualization. All tools have been ported to Python 3.

In this article, the server workflow, functionalities and performance of MDM-TASK-web are described using example data. We also includea comparison of functionality against four other servers (NAPS [1], ANCA [2], RIP-MD [3] and MDN [4]).

## 2. Materials and methods

### 2.1. Workflow for MDM-TASK-web

MDM-TASK-web is a single-page web application powered by the Django web framework [24] and a MySQL database. It relies on the Bootstrap [25] framework and the Knockout.js [26] library for a dynamic and responsive front-end, while jobs are handled by the Job Management System (JMS) [27]. The workflow used by MDM-TASK-web is shown in Figure 1.

**Figure 1:**
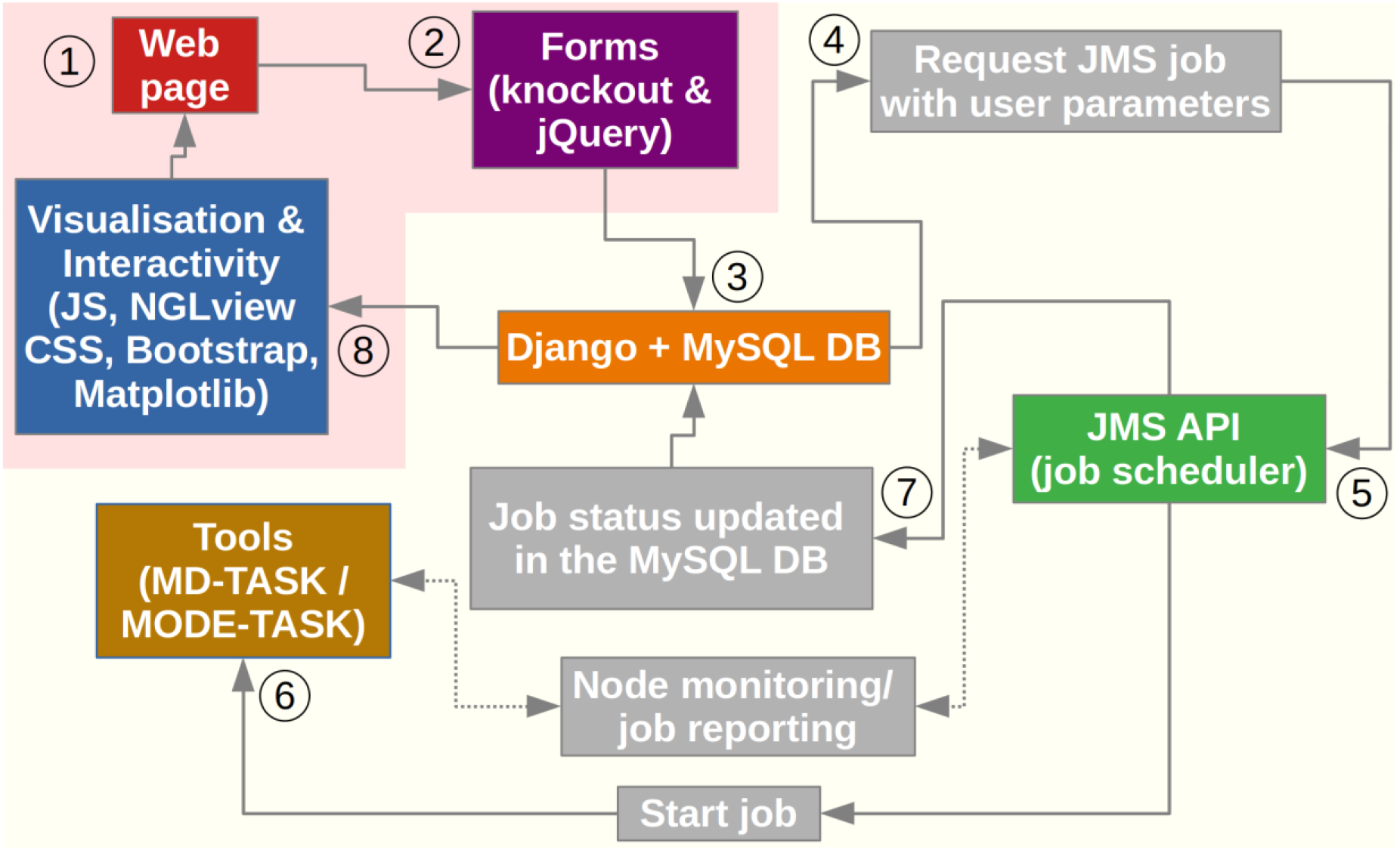
Workflow for MDM-TASK-web. The flow of execution is numbered, starting with user inputs, and ends with the visualization stage, along the unidirectional arrows. Double-sided arrows denote the two-way communication handled by JMS. Internal processes are shown in gray boxes. Front-end and back-end functionalities are highlighted with a light red and yellowish background, respectively.

Inputs from the web page are handled using jQuery and Knockout.js, and job states are synchronized with a MySQL database, which is itself managed by the Django framework. The JMS API monitors job states and compute node availability in order to schedule job submission. Job status (failure or success) is reported to JMS, which then updates the job record in the MySQL database. Depending on the tool and the job state, the relevant outputs are directly displayed using the 2D figures generated by the MDM-TASK-web Python scripts, or visualized in 3D using the NGL Viewer (version 2) [28] web application.

### 2.2. 3D visualization and tool documentation

Calculated metrics (such as correlations from perturbation response scanning (PRS), network centrality values and normal modes) are mapped onto a user-provided protein structure to facilitate interpretation where possible, via the NGL Viewer. A dedicated viewer is also available for generic trajectory visualization. Tool documentation is implemented via tool tips, collapsible buttons and demonstration pages.

### 2.3. Trajectory management

Transfer and storage of MD data can be a challenge for web servers as working with these large files can be offset by bandwidth and storage limitations. MDM-TASK-web can re-use trajectories and suggests preliminary solvent removal from the trajectory and topology files. A coarse-graining tool (https://github.com/oliserand/MD-TASK-prep) (compatible with MDTraj [29], PYTRAJ [30], MDAnalysis [31], GROMACS [32], VMD [33] and CPPTRAJ [34]) can also be used to reduce trajectory sizes by retaining only C_α_ and C_β_ atoms. Topology and trajectory files can be provided as URLs, and uploaded trajectories are re-usable. Together, these features minimize bandwidth usage and facilitate the processing of remotely simulated data without the need for specialized hardware onsite. While user data is privately stored on the server, trajectory data is removed after 30 days.

### 2.4. MD-TASK functionality

MD-TASK provides tools for performing DRN analysis, weighted contact network calculations, dynamic cross correlation (DCC) and perturbation response scanning (PRS) calculations. These are detailed in the sub-sections below.

#### 2.4.1 DRN and RIN calculations

The previous implementation of DRN analysis has been upgraded to eight centrality metrics, and now include betweenness centrality (BC), average shortest path lengths (*L*), closeness centrality (CC), eccentricity (ECC), degree centrality (DC), eigencentrality (EC), PageRank (PR) and Katz (KC) centrality that are each computed for each frame. These are then summarized for each residue as a mean, median or a standard deviation. The tool has also been adapted for single protein conformations (RIN centrality metrics). Details of DRN calculations are given in Table 1:

**Table 1:**
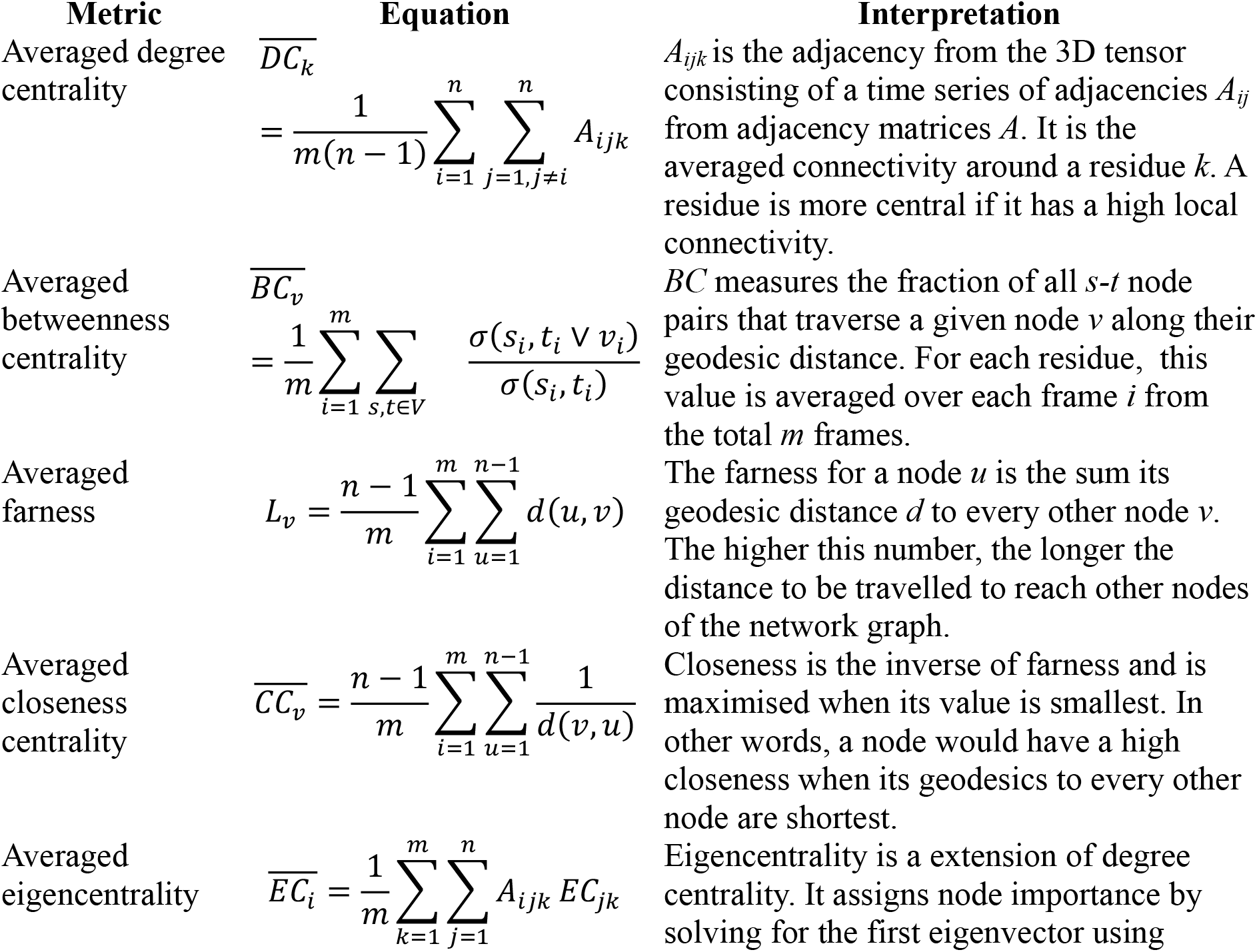
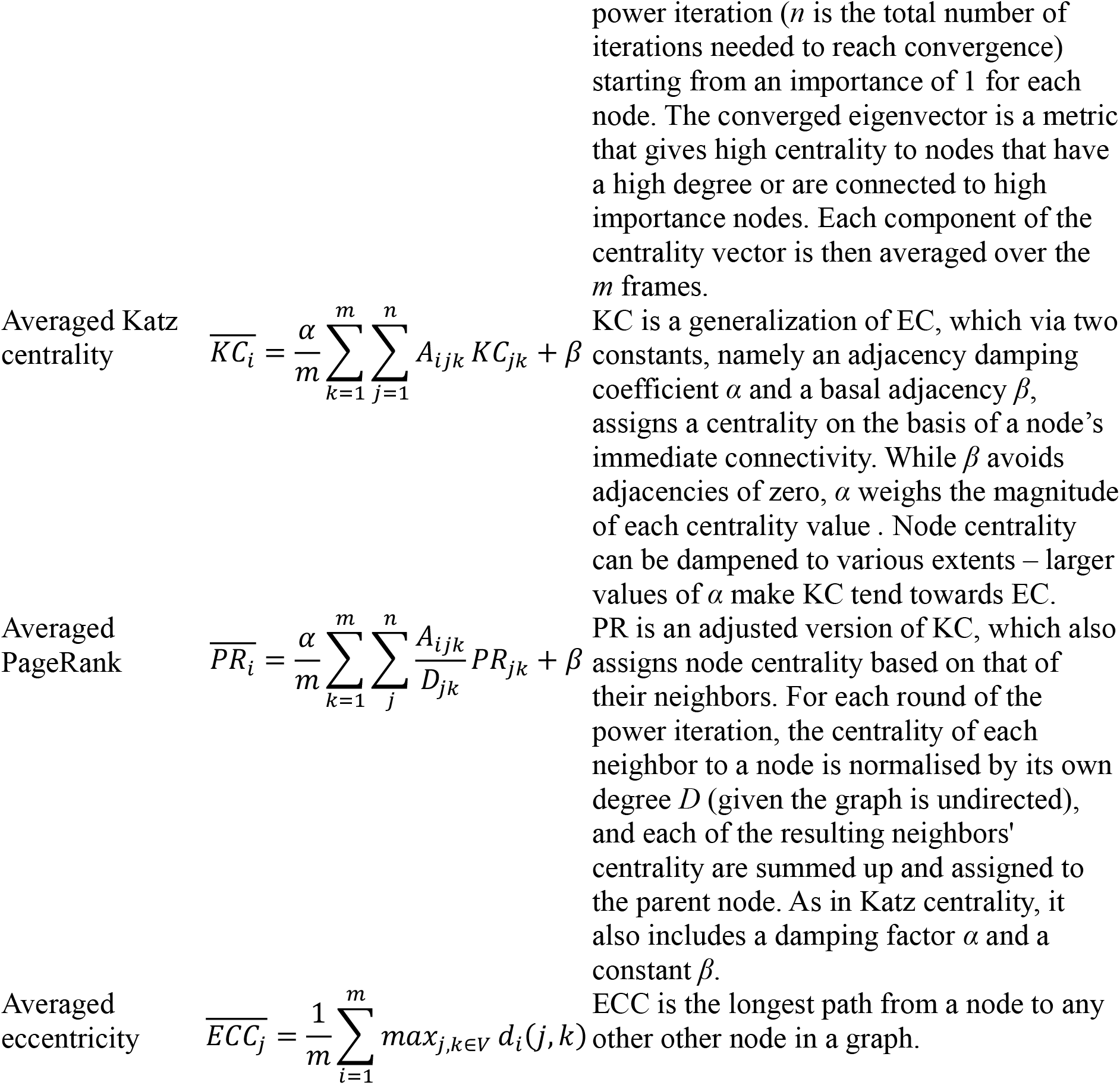
DRN metrics and their interpretations (adapted from [35])

In DRN analysis, the selected network centrality metric [35] is computed for each MD frame, and the residue centrality values are aggregated as medians or time-averages. The mapped 3D structures can be directly visualized and compared in MDM-TASK-web. Mappings are also saved in the PDBx/mmCIF format, with each DRN metric stored in the B-factor field. CSV files of the DRN metrics are also generated.

#### 2.4.2 Weighted residue contact network calculations

The original R implementation from MD-TASK was ported to Python 3, with the ability to aggregate multiple residue contacts to produce a heat map. The Ego network is calculated at one selected residue locus - this would generally be a common position, or a mutation position across several related proteins.

#### 2.4.3 Perturbation response scanning

The PRS back-end script is unchanged from that of MD-TASK; however the web interface simplifies the required inputs to requires only a trajectory, an initial conformation (PDB-formatted topology file) and a target conformation (also in PDB format) to generate an interactive 3D map of residue correlations.

#### 2.4.4 Dynamic cross-correlation

DCC, which shows the correlated residue motions, has been upgraded to work with protein complexes containing non-protein atoms.

#### 2.4.5 Communication propensity

Pairwise communication propensity is computed as the mean-square fluctuation of the inter-residue distance, using C. It relies on the fact that intra-protein signal transduction events are directly related to the distance fluctuations of communicating atoms [22,23]. Low CP values correspond to more efficient (faster) communication compared to larger values.

### 2.5. MODE-TASK functionality

MODE-TASK enables the calculation of protein ED, and the estimation of normal modes both from coarse-grained static proteins under the assumptions of the elastic network model and MD trajectories.

#### 2.5.1 Normal mode calculations from static proteins and their MD simulations

In ANM, the user is guided from the initial (optional) coarse-graining step, to solving and visualizing the normal modes. Arrows are colored by chain. Mean square fluctuations from all modes and from the first 20 non-trivial modes are separately displayed. NMA can now be computed from MD simulations as well, to represent the first dominant motion. For this calculation, the transpose of the reshaped Cartesian coordinate tensor is mean centered and dotted with its transpose to obtain the covariance matrix of dimension *3N x 3N* (where *N* is the number of residues). This matrix is then diagonalised by eigen decomposition to retrieve the principal components, in descending order of eigenvalue. The percentage of explained variance is then displayed for the first 50 modes, together with the 3D mapping of the NMA using the protein topology file. A multi-PDB file is also produced to show the mode animation. Two parameters (the *ignc* and *ignn* parameters) control the number of C- and N-terminus residues to ignore from the covariance matrix. These were included as a means to decrease possible technical variation from the termini. In our experience with protein MD simulations, we have often observed relatively high levels of fluctuation at the C-terminus.

#### 2.5.2 Essential dynamics, with improvements for comparing pairs of protein simulations

Essential dynamics tools (multidimensional scaling, standard PCA, internal PCA and t-SNE) from MODE-TASK are integrated with basic default options. A new tool, which performs comparative ED aligns one trajectory to a reference trajectory before performing a single decomposition to lay out all conformations on a common set of principal axes, such that the percentage of explained variance is the one shared by both trajectories. Comparative ED features automated conformation extraction from lowest energy basins and applies k-means to sample centroid conformations from the first 2 principal components in standard PCA. N- and C-terminal residues may be deselected before the structural alignment step to reduce unwanted noise and improve performance. Residue selection is also enabled, and is applied post global fitting of the C_α_ atoms. This is to be used only for residue and chain selections. When more than two samples are to be compared, the stand-alone script is recommended for processing all samples at once (and not as multiple paired runs), and one should also consider the percentage of explained variance captured by the principal axes. Despite its name, the comparative ED tool can also be applied to a single trajectory, in which case it will not differ from the standard PCA algorithm, if all atoms are selected.

## 3. Results and discussion

For a demonstration of use cases of MDM-TASK-web, trajectory-based tools are evaluated using mutants of the dimeric HIV-1 protease (198 residues) [36], while the enterovirus 71 capsid pentamer (PDB ID: 3VBS; 842 residues) is used for demonstrating the calculation of the anisotropic network model, which is based from a single protein conformation. In most cases, a topology and an MD trajectory are needed. In these cases, the step size parameter controls the frame sampling rate and speed of calculations, which are inversely related. Another factor that can play a potentially important role in the stability of the various trajectory-based calculations is the equilibration state of the protein. Residual effects from prior temperature and/or pressure equilibration will to some degree influence any of the aggregated metrics, and one may benefit from removing such artifacts.

### 3.1. The MDM-TASK-web interface

The web server tools each have a section where the job inputs are specified, as shown in Figure 2, for the standard PCA tool. All the tools are listed on the top menu and they generally require an MDTraj-compatible topology file and its trajectory (e.g. from GROMACS or other MD simulation tools), unless specified otherwise. In addition to the documentation sources shown in Figure 2, those of MD-TASK and MODE-TASK are also embedded in the “USER HELP” section, for further reference. The demonstration page further gives a use-case example of each tool.

**Figure 2:**
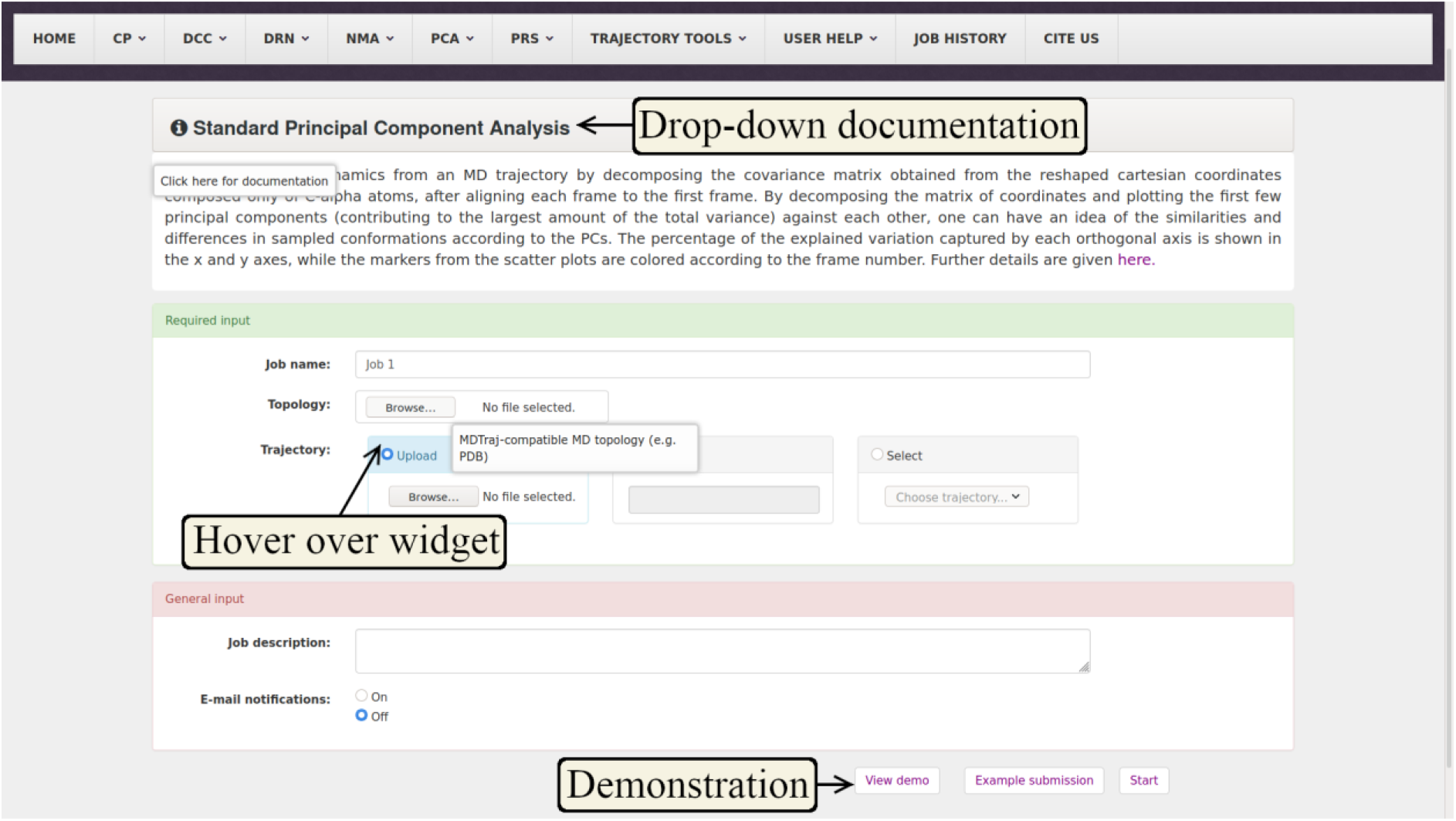
Example of the MDM-TASK-web interface, showing the embedded sources of documentation. Documentation is mainly embedded within each input page via a drop-down button, hoverable tool tips and a demonstration page.

### 3.2. Dynamic residue network & residue interaction network centrality calculations

Results from DRN centrality calculations are mapped using the user-supplied topology, as shown in Figure 3. The same back-end tool is also able to compute centrality calculations from single protein conformations when it is provided with the same file for both the topology and trajectory (under the RIN section). The paired visualizer enables the comparison of related calculations, for instance averaged EC is compared to a single conformation EC for the same protease in Figure 3. It can be seen from the averaged network centrality that the floor of central cavity (the catalytic aspartic acid) has a very high average eigencentrality, indicating that it is likely to be connected to other well-connected residues of the active site. This is mainly due to networks of H-bonding interactions and usage of the “fireman’s grip” in the protease [37]. However, there are some slight differences in centrality values when compared to those computed from the static structure. This indicates that while the static structure is faster to compute and more accessible to proteins that have not undergone MD simulations, it may be limited by the reduced sampling, especially due to the approximations taken by coarse-graining. It is also possible to use median residue centrality values, which if significantly different from the default DRN metric type (averaged centrality), would indicate the presence of skewed or multi-modal distributions, and would be the more appropriate metric to use in such cases.

**Figure 3:**
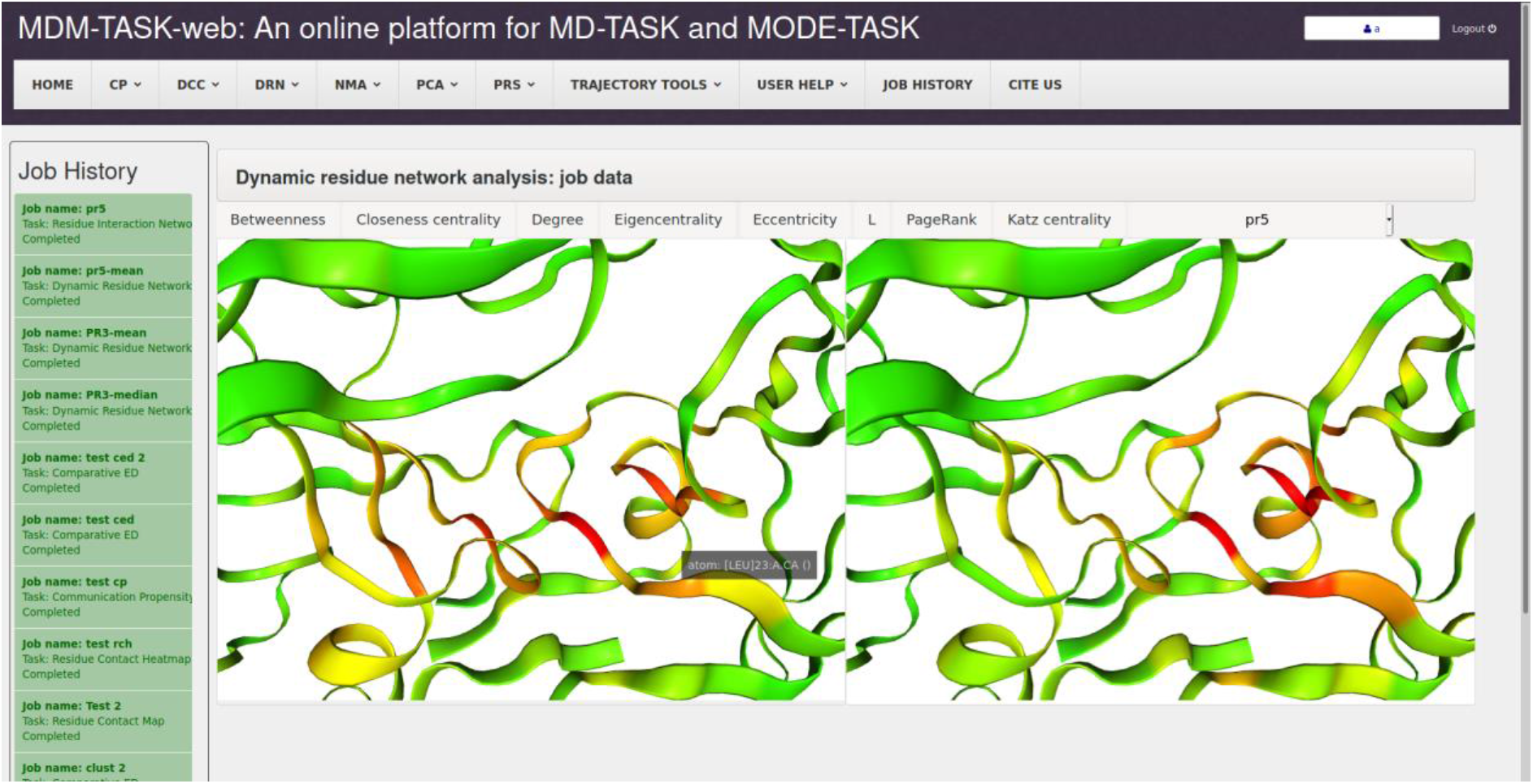
Averaged EC computed from MD frames (right side) versus EC computed from a single conformation (left side), shown in the interactive 3D interface of the MDM-TASK-web result page. Previously specified centrality metrics in the input form are shown as tabs, while the job history is shown on the left.

Further, in PyMOL [38] it is straightforward to use the PDBx/mmCIF-formatted files to visualize and compare related metrics for multiple related proteins using the *“spectrum”* command (with the B-factor values) combined with the *“set grid_mode”* command. The computed centrality metrics are all saved in CSV file, and can thus be used for customized analyses by the users.

### 3.3. Weighted residue contact network and heat maps

Weighted residue contact maps are a helpful functionality for examining local contact frequencies, for instance following averaged network centrality calculations. Figure 4 (a) shows the residue contact frequencies around GLN18 in an HIV protease mutant. Each map is associated with a file of weighted edges that can be aggregated and summarized using the contact heat map tool, for larger scale comparisons of a given locus across several protein samples, as shown in Figure 4 (b).

**Figure 4:**
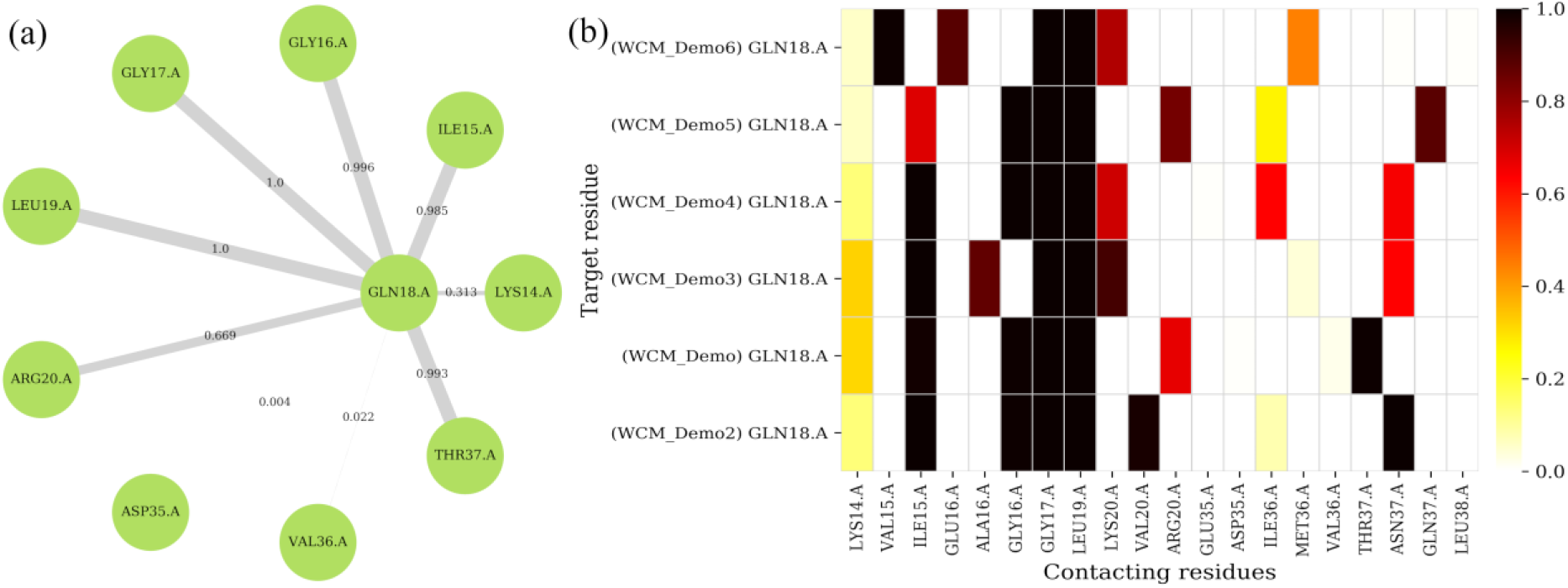
Estimating contact frequencies around a single residue in (a) a single protease, and (b) multiple HIV protease mutants, at the same locus.

### 3.4. Communication propensity

By computing the CP metric, one is able to investigate residue pairs that are the more or less likely to maintained their distances. As an example of the interpretation of the CP metric, the topology and trajectory of an HIV protease mutant (Figure 5) were used, with default parameters for the tool. From the figure, it can be seen some residue pairs display relatively higher distance variations [for e.g. between residue index pairs (37, 50), (37, 135), and (40, 91)], indicating reduced stability between these loci. This can be a preliminary investigation before proceeding to more detailed analyses.

**Figure 5:**
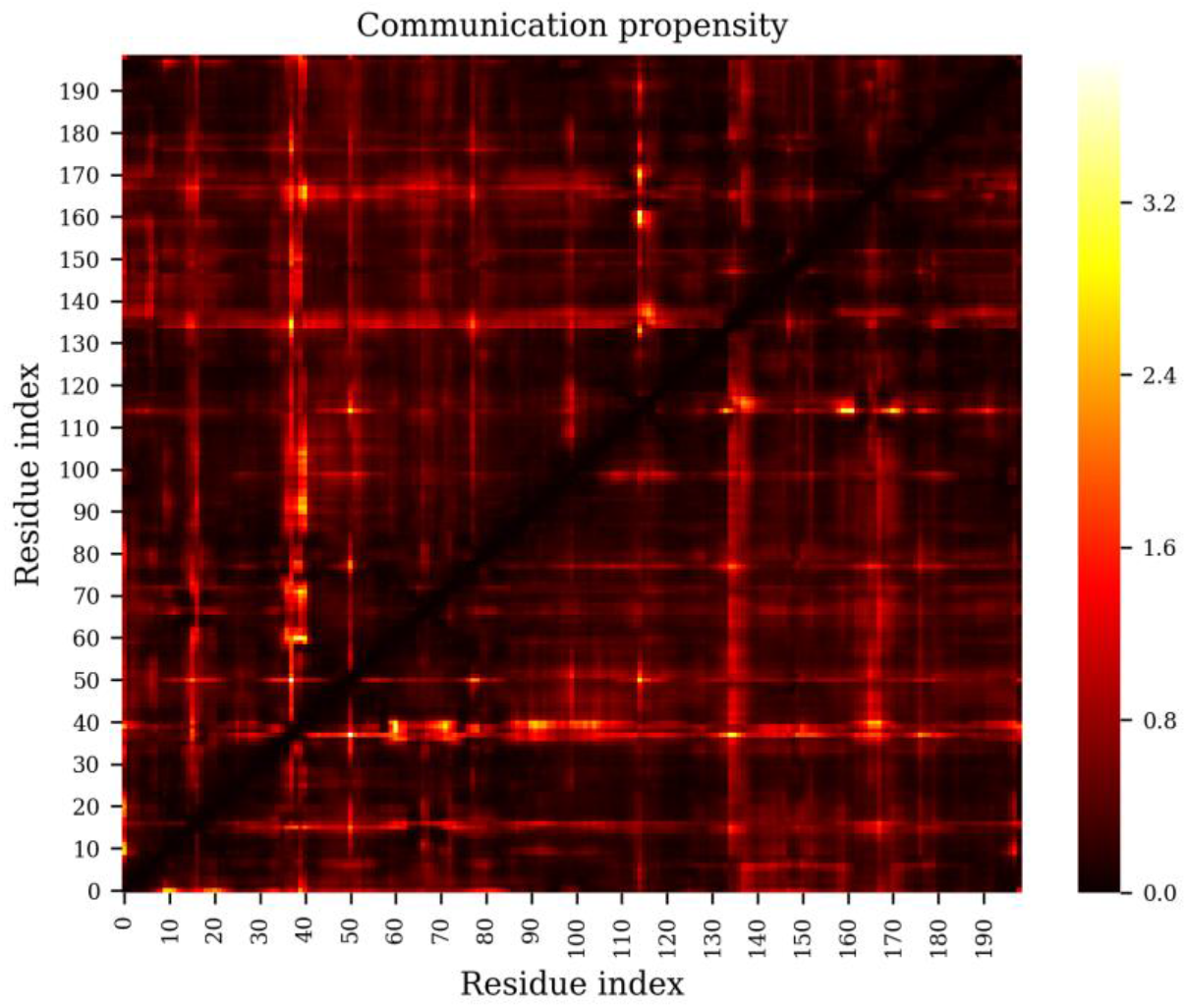
The coordination propensity calculation shows the variance in the distance between residue pairs in an HIV protease mutant.

**Figure 6:**
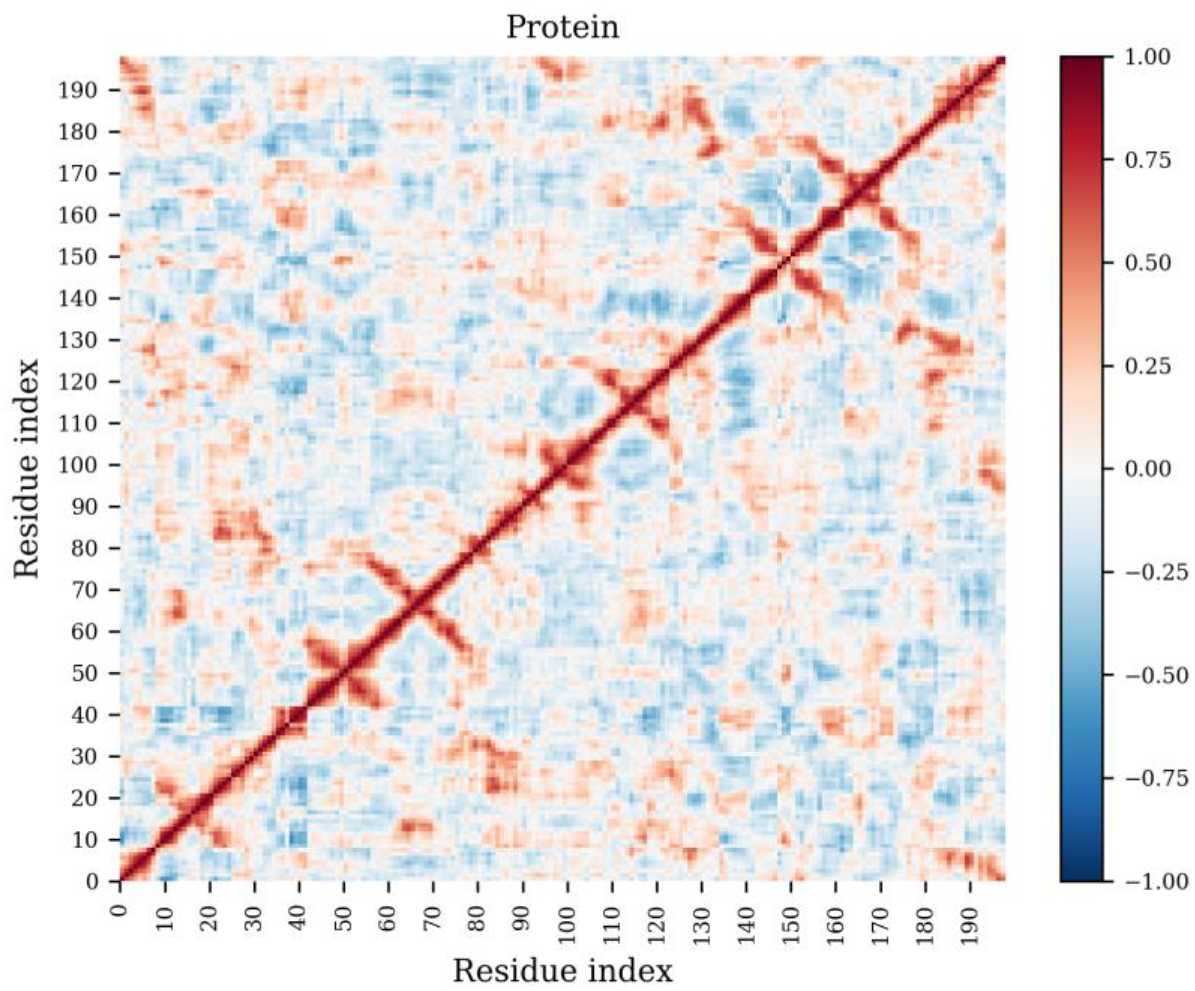
Pairwise residue correlations from an MD simulation of an HIV protease mutant. Anti-correlated and correlated movements are denoted by negative and positive DCC values, respectively, in the range [−1, 1], while uncorrelated motion has a value of zero.

### 3.5. Dynamic Cross-correlation

The DCC algorithm is unchanged from that of MD-TASK, with the exception that it is now faster and supports additional atom types. In the current example, a mutant HIV protease MD simulation was used. From the DCC heat map, one can inspect the trend in the movement of residue pairs – these can trend together, apart or be independent. Such analyses often detect protein segments that are functionally-related. By a judicious choice of atom type(s) (and/or trajectory data) one can for instance investigate protein/nucleic acid complexes using a comma-separated list “CA,P”.

### 3.6. Normal Mode Analysis (ANM and NMA from MD)

Normal mode analysis can be done using either a single PDB file or multiple protein conformations from an equilibrated MD trajectory. Single conformation NMA is done according to the anisotropic network model, as implemented in MODE-TASK, while eigen decomposition of the covariance matrix is used for MD data. In the case of the ANM, a coarse-graining level of four, C_β_ atoms and a cut-off distance of 24 were chosen to obtain six leading zero eigenvalues (displayed in the web server NMA workflow), corresponding to the trivial modes. By default mode 7 (1^st^ non-trivial) is displayed, as shown in Figure 7 (a), but other modes can also viewed by cycling through. From the ANM, we observe rotational motions within the enterovirus 71 capsid pentamer.

**Figure 7:**
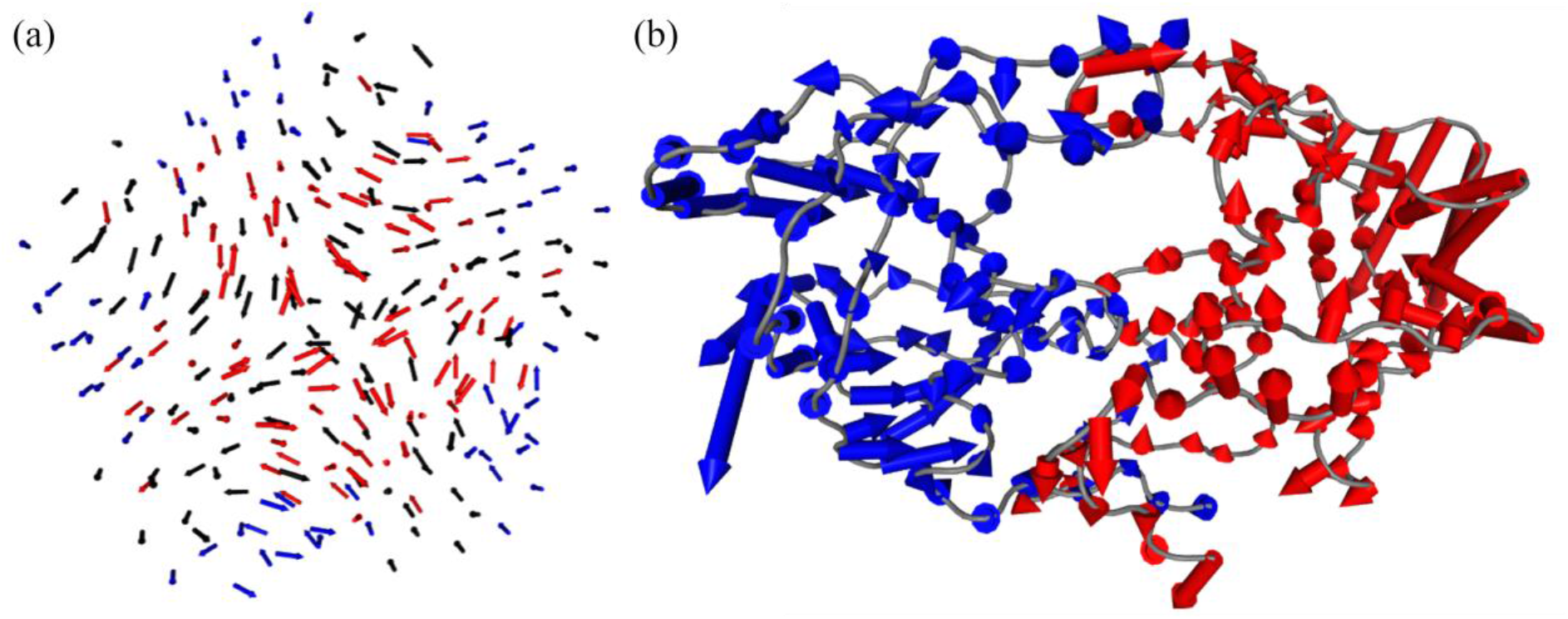
Normal mode analysis using (a) the anisotropic network model obtained from a static viral capsid pentamer, and (b) the MD covariance matrix of an HIV protease mutant. In each case, each arrow is colored by its parent chain. The arrow at each residue denotes both the extent of motion and direction with respect to each othe residue.

NMA was then computed from the MD trajectory of an HIV protease mutant (Figure 7 (b)). The decomposition differs from that used in essential dynamics in the definition of the variables. While MD frames are the variables in ED, protein residues are chosen as variables in NMA. From the first dominant motion, we observe a relatively higher amount of motion coming from the outer lateral portion of the fulcrum of the protease; internal motions are significantly smaller, with the flaps traveling only slightly more. The relative extents of motion within each chain also reveals a certain extent of protease chain asymmetry. While the back-end script can represent all modes, only one mode is represented in the web server at the moment – others may be shown in a future update.

### 3.7. Essential dynamics

ED is demonstrated via the new comparative ED tool using two HIV protease mutants that were each simulated by MD. In addition to the trajectories and topology, residues 1-31 in chains A and B were selected as example using the MDTraj syntax *‘((resid 0 to 30) and chainid 0) or ((resid 99 to 129) and chainid 1)’;* the number of clusters was set to 3 as the expected number of high probability density regions. The *ignn* and *ignc* parameters were set to their default values of 0 and 3, respectively, to ignore the three C-terminal residues, in each chain from the alignment stage onward. By default, the highest probability density conformations (in blue) are extracted from the centroids of the highest contour level using the k-nearest neighbor algorithm (where *k*=1) for the points within that level, while the kmeans algorithm is independently used to estimate other *n* possible levels of interest as specified from the number of clusters. In the example, conformations with the highest probability densities are represented by the conformations observed at time *t = 1148 ps* and *t = 1730 ps*, respectively for the first (Figure 8(a)) and second protease (Figure 8(b)) samples. In the first case, k-means detects another mode at *t = 484 ps*. The k-means prediction is partly stochastic but this effect is mitigated by being internally parameterized with a large number of iterations (1000) and multiple initializations (50) with different seeds.

**Figure 8:**
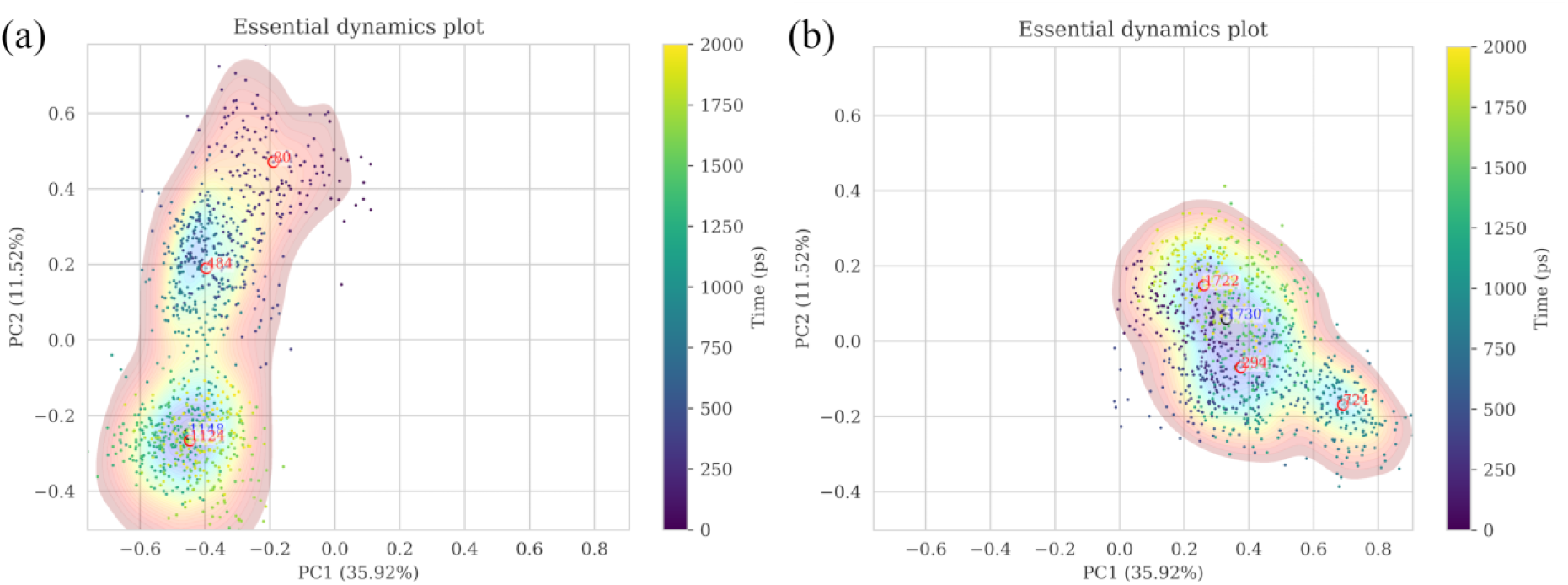
Representations of conformational sampling from independent MD simulations of two mutant HIV protease in the same eigen subspace, using comparative essential dynamics. Dots correspond to individual protein conformations (defined by a selection) and are colored by the time of sampling. The kernel density contour plots [colored from blue (lowest density) through yellow to red (highest density)] only serve as a visual guide for the energy surface, and are independently scaled, based on the respective samples. The red and blue labels are results of two separate methods for extracting conformations found at the estimated energy minima.

### 3.8. Perturbation response scanning

The interface for PRS calculation is demonstrated using a closed-conformation HIV protease mutant as starting conformation (which is also the topology file), its corresponding MD trajectory and a target conformation of the protease in an opened conformation. Applying the residue perturbation algorithm will seek for possible trigger residues that are associated with the observed conformational change of the protease flaps from a closed to an opened state. The correlations are mapped on the starting topology, as shown in Figure 9, and are also written to a text file.

**Figure 9:**
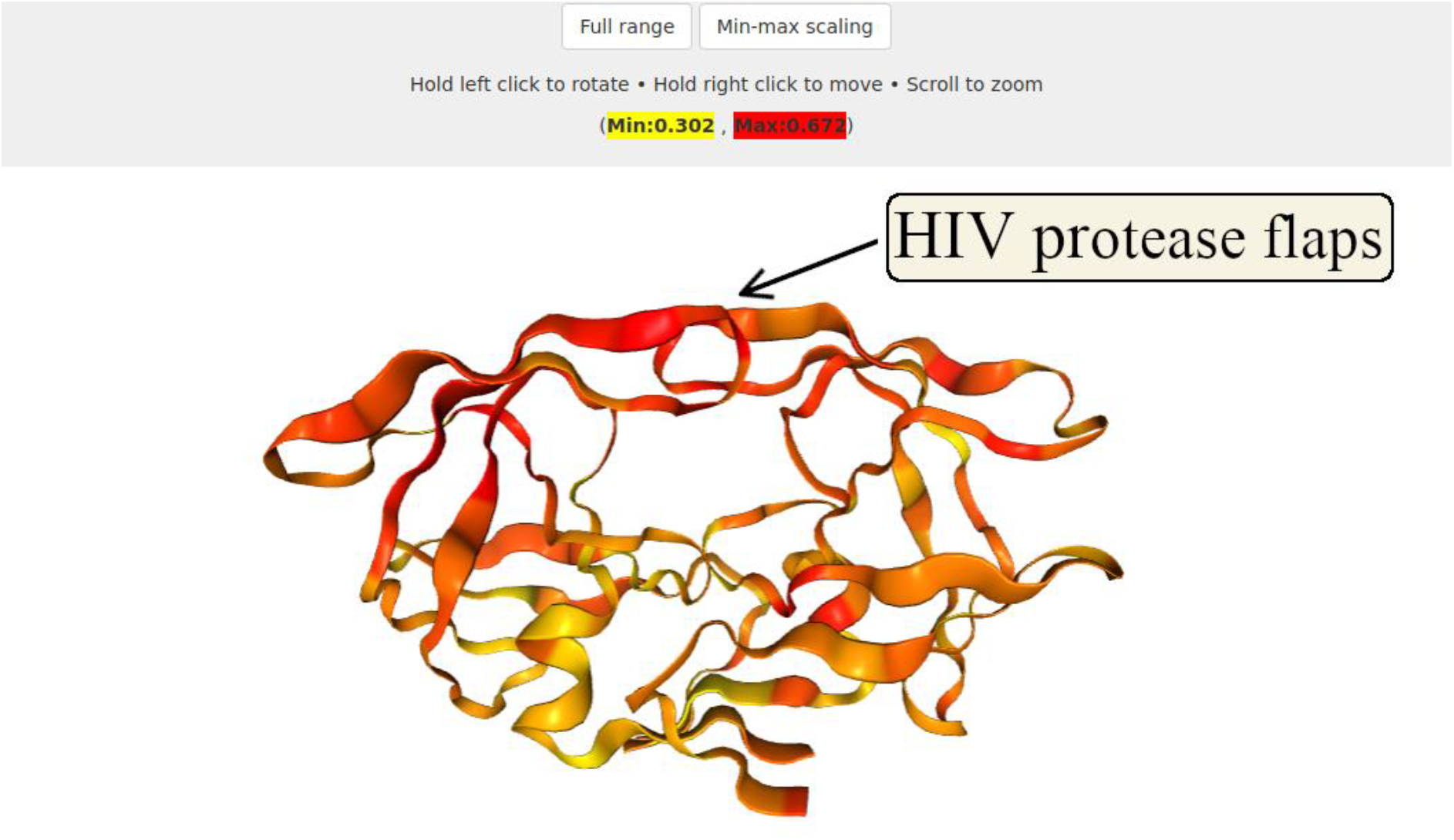
Application of PRS to scan for residues associated with the conformational shift in a closed to an opened flap conformation in HIV protease.The is left in the range [0, 1] by default, but can be scaled to the range of the observed correlations by clicking on the “Min-max scaling” option when signals have a narrower range.

## 4. Comparing MDM-TASK-web to other existing servers for MD / protein analysis

In Table 2, we show enumerate the functionalities of MDM-TASK-web and compare them against those of four existing web servers doing similar tasks.

**Table 2:**
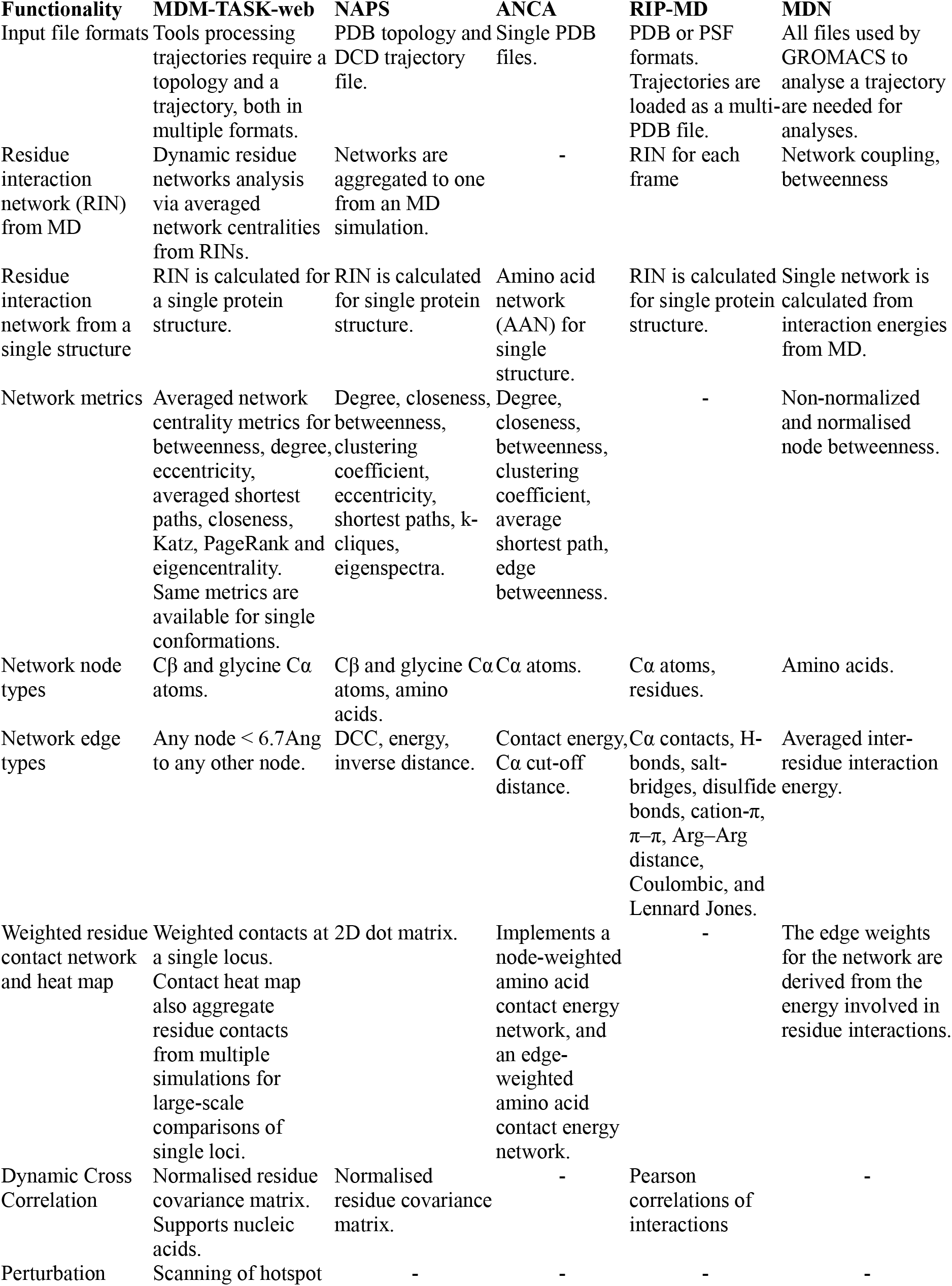
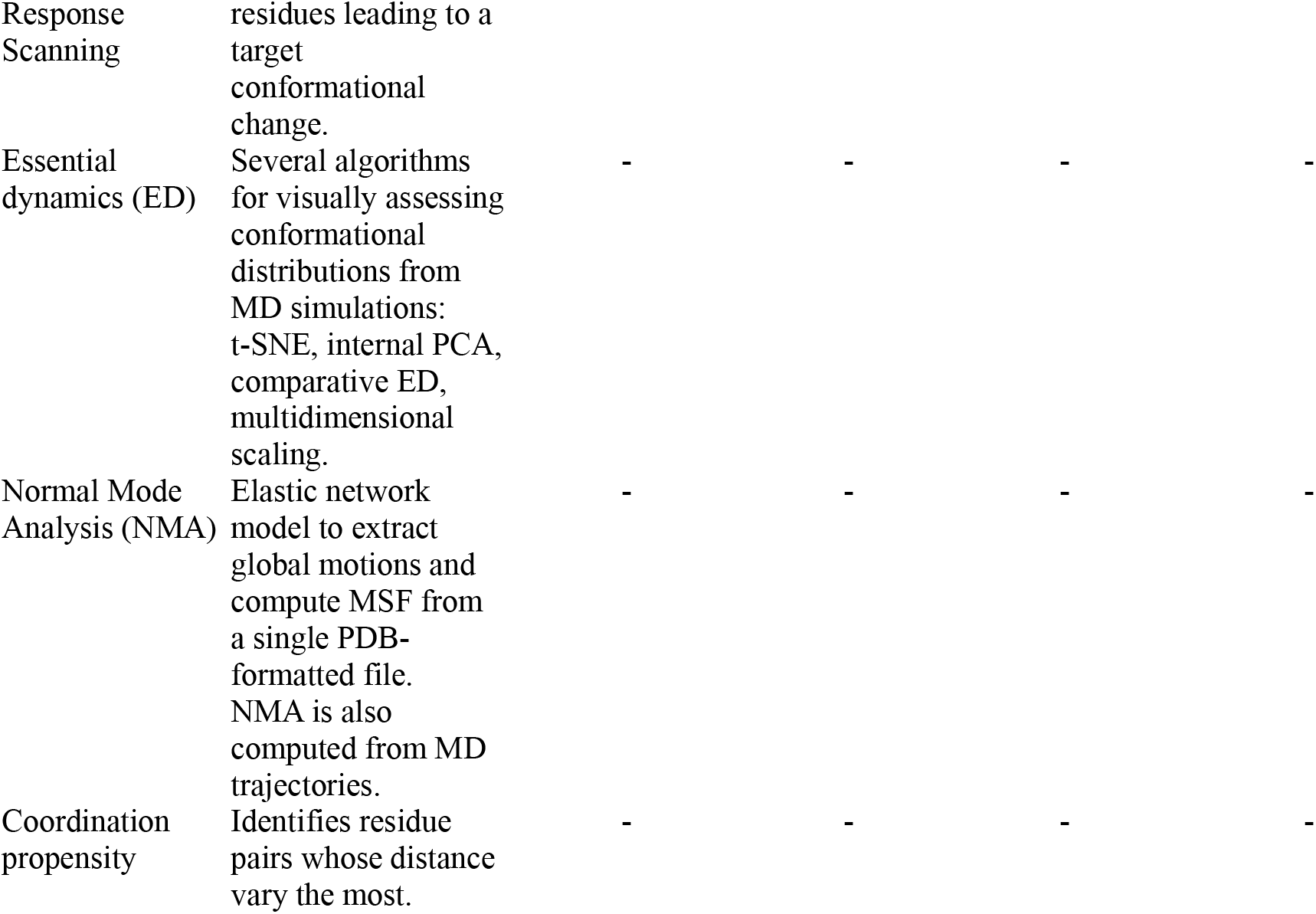
Features of MDM-TASK-web and other servers used to analyse static and dynamic protein structures

## 5. Performance

HIV protease was used to test the server-side run time of each tool in triplicate (Table 3), except for ANM where a capsid pentamer was used. Test data can be found at https://github.com/oliserand/MD-TASK-prep/example_data.

**Table 3:**
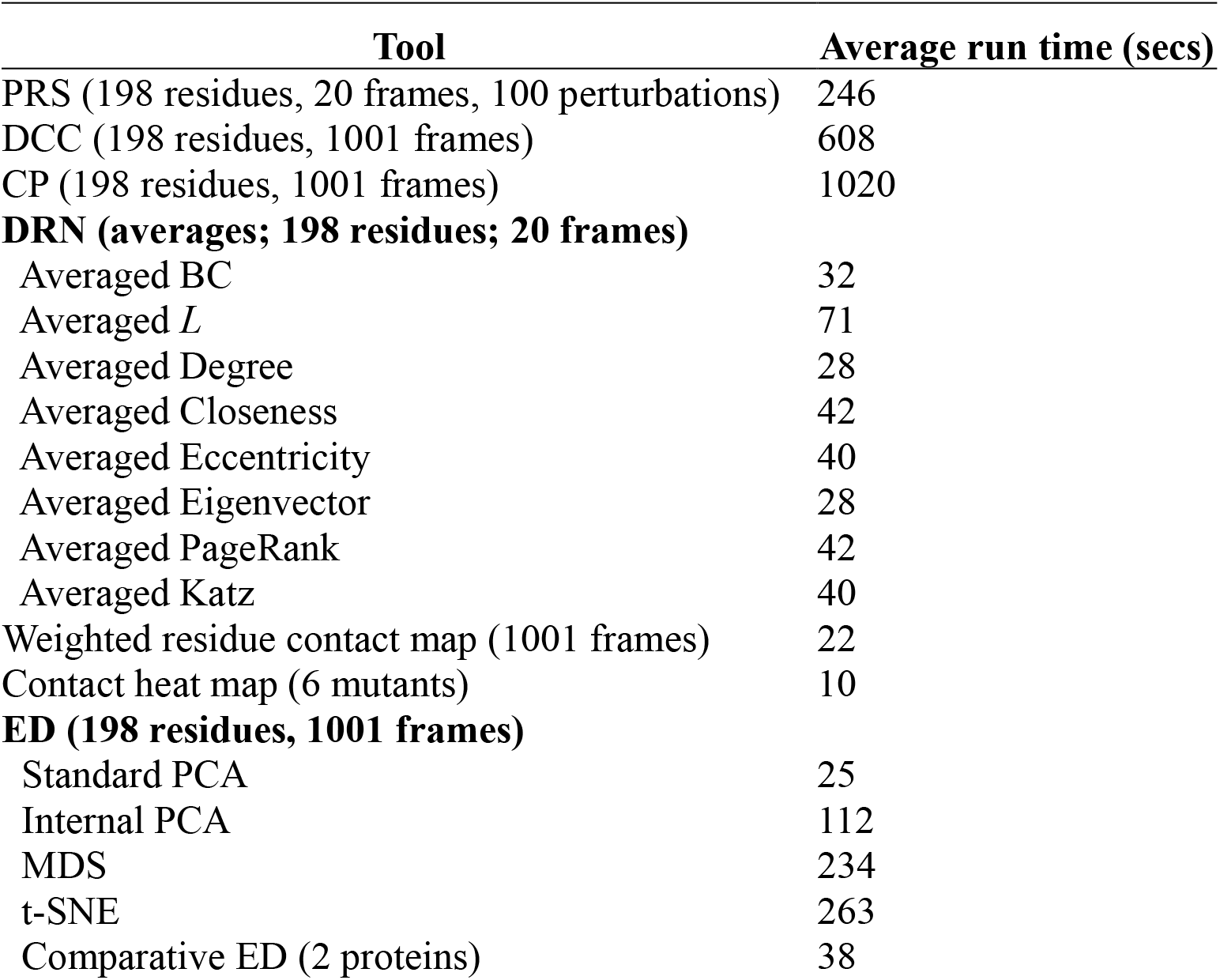
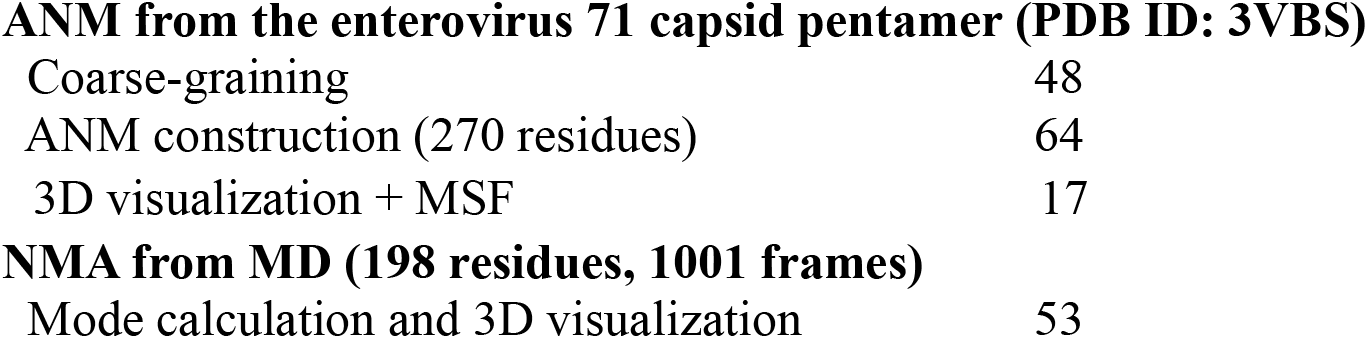
Tool performance evaluations

| Tool | Average run time (secs) |
| --- | --- |
| PRS (198 residues, 20 frames, 100 perturbations) | 246 |
| DCC (198 residues, 1001 frames) | 608 |
| CP (198 residues, 1001 frames) | 1020 |
| **DRN (averages; 198 residues; 20 frames)** |  |
| Averaged BC | 32 |
| Averaged $L$ | 71 |
| Averaged Degree | 28 |
| Averaged Closeness | 42 |
| Averaged Eccentricity | 40 |
| Averaged Eigenvector | 28 |
| Averaged PageRank | 42 |
| Averaged Katz | 40 |
| Weighted residue contact map (1001 frames) | 22 |
| Contact heat map (6 mutants) | 10 |
| **ED (198 residues, 1001 frames)** |  |
| Standard PCA | 25 |
| Internal PCA | 112 |
| MDS | 234 |
| t-SNE | 263 |
| Comparative ED (2 proteins) | 38 |
| **ANM from the enterovirus 71 capsid pentamer (PDB ID: 3VBS)** |  |
| Coarse-graining | 48 |
| ANM construction (270 residues) | 64 |
| 3D visualization + MSF | 17 |
| **NMA from MD (198 residues, 1001 frames)** |  |
| Mode calculation and 3D visualization | 53 |

## 6. Conclusion

MDM-TASK-web is a user-friendly web server for performing various types of calculations aimed at obtaining different types of insights from both static and dynamic protein data sets. By providing access to these tools in this manner also makes it available to more researchers studying proteins dynamics, without spending too much time and resources on setting up specialized hardware and software environments. The possibility of coarse-graining facilitates data transfer over the web, and tremendously reduces the data storage footprint required for the calculations, making it more likely to be accessible for further analysis without requiring significant additional storage hardware. The novel algorithms and updates to both MD-TASK and MODE-TASK enhance the capacities of each of the tool suites.

## 8. Author contributions

Ö.T.B., Conceptualization; O.S.A., Formal analysis; Ö.T.B., Funding acquisition; O.S.A., and Ö.T.B., Methodology; O.S.A., Software development; O.S.A., M.G., Web server; O.S.A., and Ö.T.B., Writing—review and editing. All authors have read and agreed to the published version of the manuscript.

## 9. Acknowledgements

We thank the Centre for High Performance Computing (CHPC); Dr D. Penkler for the CP script, and the RUBi members for their valuable suggestions.

